# MACA: Marker-based automatic cell-type annotation for single cell expression data

**DOI:** 10.1101/2021.10.25.465734

**Authors:** Yang Xu, Simon J. Baumgart, Christian Stegmann, Sikander Hayat

**Affiliations:** Bayer-Broad Joint Precision Cardiology Lab, Cambridge, MA, USA; UT-ORNL Graduate School of Genome Science and Technology, The University of Tennessee, Knoxville, TN, USA; Novo Nordisk, Data Mining and Bioinformatics, Copenhagen, Denmark; Institute of Experimental Medicine and Systems Biology, RWTH Aachen University, Aachen, Germany

## Abstract

**Summary:** Accurately identifying cell-types is a critical step in single-cell sequencing analyses. Here, we present marker-based automatic cell-type annotation (MACA), a new tool for annotating single-cell transcriptomics datasets. We developed MACA by testing 4 cell-type scoring methods with 2 public cell-marker databases as reference in 6 single-cell studies. MACA compares favorably to 4 existing marker-based cell-type annotation methods in terms of accuracy and speed. We show that MACA can annotate a large single-nuclei RNA-seq study in minutes on human hearts with ~290k cells. MACA scales easily to large datasets and can broadly help experts to annotate cell types in single-cell transcriptomics datasets, and we envision MACA provides a new opportunity for integration and standardization of cell-type annotation across multiple datasets.

**Availability and implementation:** MACA is written in python and released under GNU General Public License v3.0. The source code is available at https://github.com/ImXman/MACA.

**Contact:** Yang Xu, Sikander Hayat

## 1 Introduction

Identifying constituent cell-types in a single-cell dataset is fundamental to understand the underlying biology of the system. Many computational methods have been proposed to automatically label cells, and a benchmark study shows that a standard Support Vector Machine(SVM) classifier outperforms most other sophisticated supervised methods and can achieve high accuracy in cell-type assignment (Abdelaal, et al., 2019). However, due to lack of ground-truth in most single cell studies, supervised classification approaches are not feasible and may not be generalized for new single cell studies with different experimental designs. Therefore, unsupervised clustering approaches are still the predominant options for single-cell data analysis (Lähnemann, et al., 2020). Unsupervised approaches usually require human assistance in both defining clustering resolution and manual annotation of cell-types. This results in cell-type annotation being time-consuming and less reproducible due to human inference. As more single cell studies are available, summarizing markers identified in these studies to construct a marker database becomes an alternative approach for automatic cell-type annotation. For example, PanglaoDB (Franzén, et al., 2019) and CellMarker (Zhang, et al., 2019) are two marker databases that summarize markers found in numerous single cell studies and cover a broad range of major cell-types in human and mouse. Also, NeuroExpresso (Mancarci, et al., 2017) is a specialized database for brain cell-types. Taking advantage of those databases for robust cell type identification, we present MACA, a **m**arker-based **a**utomatic **c**ell-type **a**nnotation method and show how MACA automatically annotates cell-types with high speed and accuracy. We envision MACA as an aid for cell-type annotation to be used by both experts and non-experts.

## 2 MACA implementation

MACA takes as input expression profiles measured by single cell or nuclei RNA-seq experiments. MACA calculates two cell-type labels for each cell based on 1) an individual cell expression profile and 2) a collective clustering profile. From these, a final cell-type label is generated according to a normalized confusion matrix (Figure 1a). MACA first computes cell-type scores for each cell, using a scoring method based on a marker database or user-defined marker lists. The scoring method uses the raw gene count to calculate a cell-type score for each cell, according to gene markers of this cell-type. This results in converting a gene expression matrix to cell-type score matrix. Then, MACA generates a label (Label 1) for each cell by identifying the cell-type associated with the highest score. Independently, using the matrix of cell-type scores as input, the Louvain community detection algorithm (Blondel, et al., 2008) is applied to generate Label 2, which is a clustering label to which a cell belongs. Since the number of cell types is usually unknown, MACA tries clustering at greater resolution to over-cluster cells into many small but homogeneous groups.

**Figure 1.**
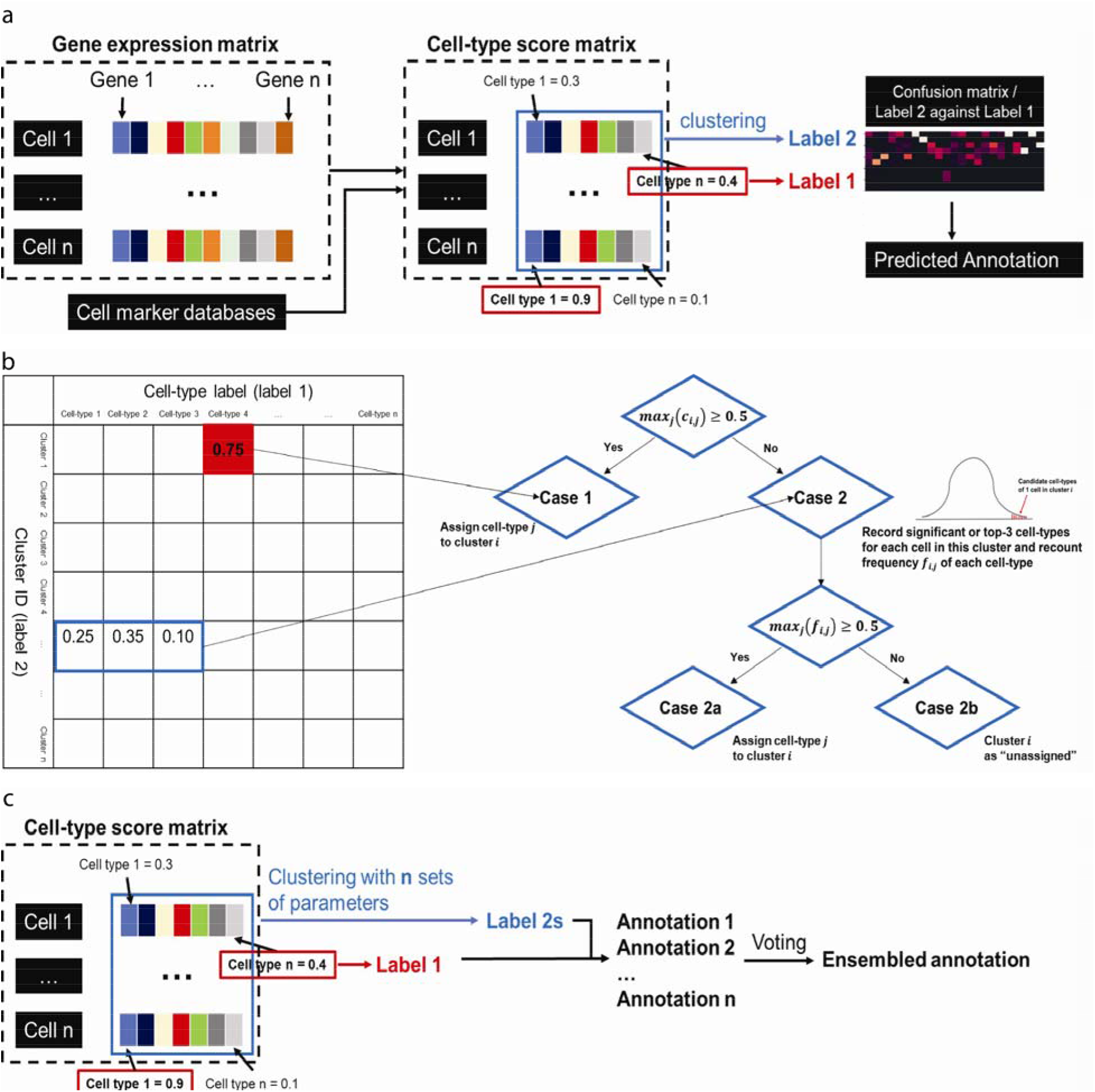
Schematic workflow of MACA. a, MACA converts gene expression matrix into cell-type score matrix based on cell marker database. MACA generates Label 1 by using max function and Label 2 by over-clustering all cells into small groups. MACA finally maps Label 2 to Label 1 via confusion matrix. b, Use of confusion matrix for cell-type annotation. c, In practical implementation, *n* sets of clustering parameters are used to generate *n* Label 2s. Mapping all Label 2s to Label 1 returns multiple annotations, and MACA ensembles these annotations by voting to generate the final cell-type prediction.

Both Label 1 and Label 2 serve complimentary functions. Label 1 is assigned on a per-cell basis which may result in incorrectly annotating many cells due to noisiness in the maximum cell-type score for each cell. This may occur when the putative cell-type feature is covered up by ambient RNAs from dominant cell-types (Pliner, et al., 2019). On the other hand, Label 2 is likely to suffer from a common problem in single cell RNA-seq clustering analysis, where cells may share the same dominant features, even though they have been clustered into different groups because of subtle differences. Additionally, results from a clustering analysis can often vary since clustering is non-deterministic. Due to its dependence on user’s decisions, mostly the choices of clustering resolution and neighborhood size.

To address these issues, MACA combines Label 1 and Label 2 to get a comprehensive cell-type annotation by mapping Label 2 to Label 1 through a normalized confusion matrix. In the confusion matrix *C*, *c*_*i,j*_ represents the number of cells that were clustered as the *i*^*th*^ cluster in Label 2 and labeled as the *j*^*th*^ cell-type in Label 1. The basic assumption of mapping Label 2 to Label 1 through a confusion matrix is that cells with the same clustering label (Label 2) should have the same cell-type label (Label 1). Ideally, if cells were identified to be in the same cluster, they should all share the same cell-type, and this cell-type has the highest score for cells in that cluster. However, in real data, this is rarely the case, as we argued above. Therefore, using a confusion matrix, we look for consensus between Label 1 and Label 2, by searching for the highest cell-type score in each cluster. Here, we compute the normalized confusion matrix *C*_*n*_ through dividing confusion matrix *C* by the size of the cluster: 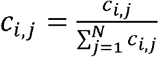, and we search for column number with the largest value for each row (Figure 1b). If *max*_*j*_(*c*_*i,j*_) ≥ 0.5, the *i*^*th*^ cluster would be assigned as the *j*^*th*^ cell-type, as more than 50% of cells in the *i*^*th*^ cluster are labeled as the *j*^*th*^ cell-type (Case 1). For cases where *max*_*j*_(*c*_*i,j*_) < 0.5, it is likely that cell identities of some cells were covered up by ambient RNAs from dominant cell-types (Case 2). Therefore, MACA records significant or at least the top-3 cell-types for each cell in the *i*^*th*^ cluster based on cell-type scores. To find significant cell-types for each cell, we get a distribution of scores of all cell-types for each cell and define those cell-types as significant if their z-scores > 3. If the number of significant cell-types is less than 3, we would keep the top-3 cell-types. Doing this can retrieve more potential cell-type labels for this cluster, and each cell will contribute at least 3 candidates into a pool of candidate cell-types for this cluster. Then, MACA calculates frequency of each candidate cell-type in this pool and assigns the *i*^*th*^ cluster as the cell-type with the highest frequency if the frequency exceeds half the size of the cluster (*max*_*j*_(*f*_*i,j*_) ≥ 0.5) (Case 2a). Otherwise, the *i*^*th*^ cluster would be labeled as “unassigned” (*max*_*j*_(*f*_*i,j*_) < 0.5) (Case 2b), which is the case that cells in this cluster do not have an agreement on which cell-types they belong to. For the choice of 0.5, we will show our examination in the next Results section. As we mentioned before, clustering-based cell-type identification largely depends on user’s choice, for example the choices of clustering resolution and neighborhood size. Therefore, the outcome may vary among different users. To have a more reproducible outcome, we cluster cells with different clustering parameters to get multiple clustering assignments (Label 2s). Repeating the procedure of mapping Label 2 to Label 1 will enable us to get an ensemble annotation through voting, and this ensemble annotation is less influenced by a single clustering choice (Figure 1c). Using ensemble approach also offers a naïve way of scoring MACA-based cell-type predictions. Users can set up a threshold to filter cells whose annotations are less consistent in outcomes of different clustering trials, and we also provide examinations in the next section to help users choose a reasonable threshold for annotation with quality. In this study, we generated clusters using Louvain method with 3 different resolutions and 3 different numbers of neighborhood, which results in 9 different clustering labels (Label 2s). After mapping these 9 Label 2s to Label 1, we generated 9 cell-type annotations. Then, we used a voting approach to get the final annotations (the highest votes from the 9 annotations). Users can also increase the number of clustering trials to have a larger voting pool for annotation ensemble or decrease the number to save computation time.

Back to converting gene expression matrix to cell-type score matrix, we collected 4 different scoring methods that were proposed to do the conversion. These scoring methods are either named by authors, or we named them after the last name of the first author. PlinerScore was a part of Garnett that was designed to annotate cell-types through supervised classification (Pliner, et al., 2019). The uniqueness of PlinerScore is the use of TF-IDF transformation to deal with specificity of a gene marker and a cutoff to deal with issue of free mRNA in single-cell RNA-seq data. AUCell comes from SENIC, which uses gene sets to quantify regulon activities of single-cell expression data (Aibar, et al., 2017). In this study, AUCell quantifies the enrichment of every cell-type as an area under the recovery curve (AUC) across the ranking of all gene markers in a particular cell. This assessment is cell-wise and is different from PlinerScore that requires transformation of the whole dataset. Both CIM and DingScore simply use the total expression of all gene markers of a particular cell-type as the cell-type score (Ding, et al., 2020; Efroni, et al., 2015). CIM normalizes the total expression by multiplying a weight that is defined as the number of expressed gene markers divided by the number of all gene markers of this cell-type. DingScore, on the other hand, normalizes the total expression of one cell-type by dividing total expression of all genes. Since some cell-types have a longer list of marker genes than others, cell-types with more marker genes in the database would have larger cell-type scores. Normalization in CIM was considered to address this issue. However, PlinerScore and DingScore were not intentionally designed to cope with unbalanced marker lists. To deal with this issue, we did a similar processing to normalization in CIM, which is dividing the score of each cell type by the number of expressed markers in that cell type. However, AUCell is a completely different approach from the other 3 scoring methods, which does not simply sum up values of marker genes for a given cell-type. So, we ran AUCell without extra processing for returned values. Meanwhile, we show that the number of expressed marker genes in both PanglaoDB and CellMarker across 6 single cell datasets tested in this study, and we found that most cell-types in PanglaoDB have expressed marker genes within 0~60, while most cell-types have less than 10 marker genes expressed in CellMarker (Supplementary Figure S1). For both PanglaoDB and CellMarker, we can conclude that cell-types with over 100 expressed marker genes are a minority.

In practice, we build MACA in the analysis pipeline of Scanpy, and MACA takes data in the format of “anndata” in Python (Wolf, et al., 2018). Expression data are preprocessed through cell and gene filtering, and transformed by LogNormlization method, the common practice in single cell analysis. Then, the user provides marker information in the form of Python dictionary, and MACA transforms gene expression matrix to cell-type score matrix. Next, annotation by MACA can be summarized into 4 steps as shown in Figure 1: 1) Louvain clustering to generate Label 2; 2) Generating Label 1 via max function; 3) Mapping Label 2 to Label 1 through normalized confusion matrix; 4) Repeating step 1 to 3 to have ensembled annotation.

## 3 Results

The key component for optimal performance of MACA is constructing cell-type scores from the gene expression matrix. We investigated 4 scoring methods that have been proposed to transform gene expression matrix to cell-type score matrix (Aibar, et al., 2017; Ding, et al., 2020; Efroni, et al., 2015; Pliner, et al., 2019), and we tested these methods with 2 public marker databases (Franzén, et al., 2019; Zhang, et al., 2019) in 6 single cell studies comprised of 3000 to 20000 cells (Baron, et al., 2016; Cui, et al., 2019; Tian, et al., 2019; Vieira Braga, et al., 2019; Wang, et al., 2020; Zheng, et al., 2017), which include 3 benchmark datasets (Supplementary Table S1) (Abdelaal, et al., 2019). To evaluate these annotation outcomes, we used Adjusted Rand Index (ARI) and Normalized Mutual Information (NMI). Both ARI and NMI are calculated by measuring similarity or agreement between our annotations and authors’ annotations. For the 3 benchmark datasets, authors’ annotations would be the ground truth label, while authors’ annotations in the other 3 datasets are at least created under careful investigation. Therefore, use of ARI and NMI, in this case, is to show how well we can reproduce authors’ outcomes. We found annotations using PlinerScore with markers in PanglaoDB have the largest agreement with authors’ annotations for all 6 datasets, in terms of both ARI and NMI (Table 1). Therefore, MACA uses PanglaoDB with PlinerScore as the main marker database and scoring method, respectively. When we define if Label 2 agrees with Label 1, we selected 0.5 as the threshold. It is out of a simple reasoning of whether the half agrees. However, it is possible to set up a less or more stringent threshold to define the consensus between Label 1 and 2. Thus, we further tested how different thresholds will affect MACA’s performance. We changed the threshold from 0.2 to 0.9 and performed our test in these 6 datasets. We expect annotations would vary, but surprisingly, MACA’s performance is quite robust to the choice of this parameter, except that we observed drops of ARI and NMI in two datasets when using 0.9 as threshold (Supplementary Table S2).

**Table 1.** Performance of MACA, CellAssign, SCINA, Cell-ID, and scCATACH in 6 scRNA-seq datasets, measured by ARI and NMI. 8 different settings of MACA include using 4 cell-type scoring methods (PlinerScore, AUCell, CIM, and DingScore) with 2 marker databases (PanglaoDB and CellMarker).

| ARI | PBMC (Zhenget al., 2017) | CellBench (Tian et al., 2019) | Pancreas (Baron et al., 2016) | Heart (Wanget al., 2020) | Heart (Cui et al., 2019) | Lung (Vieira et al., 2019) |
| --- | --- | --- | --- | --- | --- | --- |
| PanglaoDB+PlinerScore | **0.95** | **0.92** | **0.90** | **0.71** | **0.61** | **0.45** |
| PanglaoDB+AUCell | 0.04 | 0.00 | 0.78 | 0.39 | 0.47 | 0.29 |
| PanglaoDB+CIM | 0.28 | **0.65** | **0.90** | 0.27 | 0.30 | 0.33 |
| PanglaoDB+DingScore | **0.83** | **0.74** | 0.69 | 0.07 | 0.44 | 0.20 |
| CellMarker+PlinerScore | 0.38 | 0.43 | 0.27 | **0.57** | 0.13 | 0.21 |
| CellMarker+AUCell | 0.29 | 0.52 | 0.32 | 0.34 | 0.09 | 0.14 |
| CellMarker+CIM | 0.24 | 0.60 | 0.54 | **0.56** | 0.07 | 0.09 |
| CellMarker+DingScore | 0.22 | 0.55 | 0.38 | 0.37 | 0.19 | NA |
| SCINA | 0.46 | 0.63 | **0.89** | 0.13 | **0.55** | 0.31 |
| CellAssign | NA | 0.00 | **0.89** | 0.15 | **0.53** | 0.26 |
| Cell-ID | 0.50 | 0.17 | 0.57 | 0.10 | 0.49 | **0.35** |
| scCATCH (best) | **0.62** | 0.56 | 0.86 | 0.04 | 0.14 | **0.60** |
| scCATCH (average) | 0.57 | 0.40 | 0.66 | 0.04 | 0.05 | **0.35** |
| NMI | PBMC (Zhenget al., 2017) | CellBench (Tian et al., 2019) | Pancreas (Baron et al., 2016) | Heart (Wanget al., 2020) | Heart (Cui et al., 2019) | Lung (Vieira et al., 2019) |
| PanglaoDB+PlinerScore | **0.89** | **0.92** | **0.88** | **0.59** | **0.62** | **0.59** |
| PanglaoDB+AUCell | 0.09 | 0.00 | 0.79 | **0.41** | 0.50 | 0.31 |
| PanglaoDB+CIM | 0.51 | **0.80** | **0.88** | 0.30 | 0.44 | 0.40 |
| PanglaoDB+DingScore | 0.74 | **0.85** | 0.70 | 0.10 | 0.47 | 0.33 |
| CellMarker+PlinerScore | 0.44 | 0.64 | 0.57 | **0.51** | 0.32 | 0.42 |
| CellMarker+AUCell | 0.23 | 0.67 | 0.46 | 0.32 | 0.33 | 0.17 |
| CellMarker+CIM | 0.49 | 0.78 | 0.73 | **0.41** | 0.31 | 0.21 |
| CellMarker+DingScore | 0.43 | 0.73 | 0.60 | 0.34 | 0.33 | 0.08 |
| SCINA | 0.54 | 0.71 | 0.84 | 0.07 | **0.54** | 0.46 |
| CellAssign | NA | 0.06 | **0.86** | 0.08 | 0.51 | 0.49 |
| Cell-ID | 0.67 | 0.38 | 0.74 | 0.08 | **0.55** | 0.58 |
| scCATCH (best) | **0.77** | 0.70 | 0.84 | 0.05 | 0.30 | **0.73** |
| scCATCH (average) | **0.75** | 0.62 | 0.75 | 0.04 | 0.12 | **0.63** |
| # of cell-types | PBMC (Zhenget al., 2017) | CellBench (Tian et al., 2019) | Pancreas (Baron et al., 2016) | Heart (Wanget al., 2020) | Heart (Cui et al., 2019) | Lung (Vieira et al., 2019) |
| MACA | 8 | 6 | 11 | 8 | 7 | 13 |
| SCINA | 14 | 14 | 17 | 16 | 23 | 41 |
| CellAssign | NA | 9 | 17 | 18 | 24 | 31 |
| Cell-ID | 33 | 55 | 48 | 35 | 37 | 63 |
| scCATCH (best) | 9 | 5 | 10 | 3 | 3 | 16 |
| Author's annotation | 5 | 5 | 14 | 5 | 9 | 13 |

Next, we seek to compare MACA with other existing marker-based annotation tools. CellAssign and SCINA are two computational methods that have been proposed for automatic cell-type assignment (Zhang, et al., 2019; Zhang, et al., 2019). Both methods rely on statistical interference to compute the probabilities of cell types, which are time- and computation-intensive. Recently, Cell-ID was released for extraction of gene signature as well as cell-type annotation (Cortal, et al., 2021). We also noticed scCATCH and SCSA, which are both cluster-based annotation tools (Cao, et al., 2020; Shao, et al., 2020). Both scCATCH and SCSA require identifying differential marker genes for each cluster via a statistical test implemented in Seurat and then matching identified cluster markers to marker database (Butler, et al., 2018). Here, we compared MACA with CellAssign, SCINA, Cell-ID, and scCATCH using these 6 single cell studies and cell markers in PanglaoDB. We tested MACA, CellAssign, SCINA, Cell-ID, and scCATCH on a workstation with 16-core CPU and 64GB memory. MACA can finish annotation within 1 minute (cells around 3,000) and less than 2 minutes for a relatively large dataset (cells up to 20,000 cells). On the datasets used and on our computational resources, scCATCH and Cell-ID took longer than MACA to compute annotations and ranks as the second and third fastest. In our hands, SCINA took around 20-minute time to finish annotation for a large dataset, and CellAssign took the longest time to complete cell-type assignment and failed to annotate data with > 20,000 cells due to lack of memory (Supplementary Table S3). Because annotation by scCATCH needs clustering first and differential marker identification is highly affected by clustering outcome, the investigator will need to do a thorough investigation to make sure that clustering is not overdone or underestimated. In this study, we reported the highest and the averaged outcomes of scCATCH in each dataset. Comparing these results with manual annotations from the authors, we found 1) MACA labels cells had a higher consensus than CellAssign, SCINA, Cell-ID, and scCATCH, in terms of both ARI and NMI, and 2) MACA and scCATCH identify similar numbers of cell-types to author’s annotations, while the other 3 methods, especially Cell-ID, report overall more different cell-types (Table 1). The low ARIs and NMIs of CellAssign and Cell-ID can be counted as results of 1) many “unassigned” cells and 2) exceeding numbers of different cell-types over the numbers reported by authors. It is important to note that other methods compared here were run on their default parameters. In future, parameter tuning of those methods on a computer with higher memory should be carried out for a comprehensive benchmarking on many datasets. Finally, to better evaluate annotations, we used a machine learning approach to assess cell-type assignment. Training classifiers was recently proposed by (Miao, et al., 2020) to assist in finding a good clustering resolution, and we adopt this idea to evaluate our annotations. Basically, if the annotation is good enough, we can train a classifier to predict cell type using gene-expression values with high accuracy. Conversely, if there are many wrong labels, it would be hard for a classifier to make the right decision. We performed 5-fold cross-validated training, where we split one dataset into 4-fold training set and 1-fold testing set and trained a SVM classifier on the training sets and applied the classifier to predict labels for the testing set. This procedure repeats 5 times to get a mean accuracy. Instead of treating authors’ annotations as ground truth, this machine-learning evaluation provides an independent angle to judge annotation quality. Indeed, MACA achieves high concordance with authors’ reported annotations and higher mean of accuracies than other methods (Supplementary Table S4). Of note, high accuracy of SVM classifier is not equal to correctness of annotation. Meanwhile, ARI and NMI reports similarity between two annotations but cannot reflect the difference of annotation resolution. For example, MACA may return less cell-types than authors. Moreover, annotation resolution of MACA highly depends on the number of cell-types in the marker database, and it is likely that MACA cannot annotate some rare subtypes that do not show up in the marker database. Here, we used confusion matrix to show how cell-type labels by MACA are against cell-type labels by authors (Supplementary Figure S2). Take annotation of human pancreas as an example, cells annotated by MACA as “Pancreatic stellate cells” fall into 3 groups that were annotated by author as “activated stellate cells”, “quiescent stellate cells”, and “Schwann cells”, respectively. Since MACA may have a different annotation resolution from the author’s, we performed a test to show how different annotation resolutions can affect calculations of ARI and NMI. We included the human kidney (CD10-) data, which has 3 different annotation resolutions by the authors, from 5 major cell-types to 29 intermediate cell-types, and to 50 fine cell-types (Kuppe, et al., 2021). We used MACA to annotate this data and compared MACA’s annotation with these 3 annotations. We found NMI is more robust to change of annotation resolution than ARI. It also suggests that a higher ARI reflects similar resolution between MACA and author. (Supplementary Figure S3).

As we mentioned above, using ensemble approach also offers user an option to filter cells whose annotations are less consistent in outcomes of different clustering trials. However, it also causes loss of cells for downstream analysis, like cellular composition analysis. To find a good balance between having higher annotation quality and keeping most cells for downstream analysis, we tested threshold of voting from 1/9 to 9/9, where the numerator means the minimum number of votes required to keep the cell-annotation. With 1/9, all cells will be kept, with 2/9, cells with annotations with at least 2 votes will be kept, while only cells that have the same annotation across 9 clustering trials will be considered if threshold is set up as 9/9. We reported the results across 10 datasets in Supplementary Table S5, and it may provide a reference for user to choose a threshold that serves user’s need. Of note, we kept all cells in other evaluations. Particularly, all cells were used in benchmark with other methods. Here, we suggest setting up the threshold as 7/9. Next, we expect to show that annotation by MACA is applicable for most single cell RNA-seq platforms. We re-annotated PBMC data from a new study by (Ding, et al., 2020). This data consists of two biological samples from 9 platforms. We found that 1) both PBMC samples have the same major cell-types, and these 9 platforms can successfully profile them (Supplementary Figure S4a), and 2) annotation by MACA shows that all platforms profile similar cellular components for these two PBMC samples, except CEL-Seq2 (Supplementary Figure S4b). These results are largely consistent to the original report (Ding, et al., 2020). However, this PBMC data didn’t come with a ground-truth annotation, we further added the human pancreas data, which consists of 5 independent studies profiled by 4 different single-cell RNA-seq platforms (Baron, et al., 2016; Grün, et al., 2016; Lawlor, et al., 2017; Muraro, et al., 2016; Segerstolpe, et al., 2016). Annotation by MACA has 0.929 ARI and 0.908 NMI against author-reported annotation, and we also observed all major cell-types were revealed across all 4 platforms (Supplementary Figure S4c).

Finally, we applied MACA to a single-nuclei RNA-seq dataset from all 4 chambers of the human heart, comprised of ~290k nuclei (Tucker, et al., 2020). MACA could annotate each of the 4 chambers comprising of ~80K cells each in < 6 mins. Annotations by MACA have major agreement with author’s reported annotations with an average ARI and NMI of 0.63 and 0.76, respectively (Supplementary Table S6). However, we also found some disagreements exist in annotation of cells in from left and right atria. Therefore, we investigated disagreement between MACA’s and author’s annotations, and found the biggest difference stems from disagreement in assignments for neuronal cells and lymphocytes, which are both small-population cell types in this dataset (1702 neuronal cells and 1503 lymphocytes out of ~290k). We found neuronal cells weren’t revealed and author-reported lymphocytes were reported as memory T cells in MACA’s annotation (Supplementary Table S7a and b).

By default, MACA works with the list marker genes and cell-types present in PanglaoDB, but users can also input their own gene-lists. A major limitation of MACA is that it can only annotate cell-types that are pre-defined in the marker reference, but with more marker gene-sets becoming available with single-cell sequencing studies, we believe that MACA will be useful to annotate heterogeneous single-cell datasets. This points us two future directions to improve MACA. First, with more atlas studies that profile all sorts of biological systems, more refined markers for small cell populations can be defined, and MACA could reach finer annotation resolution by integrating markers from these new atlas studies. Second, weights of markers should be incorporated into the scoring method of MACA, for example marker specificity and expression strength. However, at the current stage, all markers have equal weights when they contribute to cell-type scores, and we believe that incorporating marker weights will be beneficial for accurate annotation. With a more refined marker database and cell-type scoring method, MACA would rapidly perform integrated annotation across multiple datasets, and this is very critical for downstream analyses like cellular component analysis across datasets under different conditions. In fact, we noticed that combining PlinerScore and PanglaoDB to generate new features has the advantages of correcting batch effects for integrated annotation across datasets, and we aim to extend the use of MACA to standardization of cell-type annotation across datasets in the future (see application in integrated annotation on GitHub of MACA). Finally, we conclude that MACA is a suitable tool for automatic cell-type annotation that can aid both experts and non-experts in rapid annotation of their single-cell datasets.

## Acknowledgements

We would like to thank Mark Chaffin, Stephen Fleming and other members of the Precision Cardiology Lab for providing useful feedback on the manuscript.

## Disclosures

Simon Baumgart, Christian Stegmann and Sikander Hayat are employees of Bayer US LLC (a subsidiary of Bayer AG) and may own stock in Bayer AG.

## Notes

### Competing Interest Statement

The authors have declared no competing interest.

## References

Abdelaal, T., et al. A comparison of automatic cell identification methods for single-cell RNA sequencing data. Genome biology 2019;20(1):194–194.

Aibar, S., et al. SCENIC: single-cell regulatory network inference and clustering. Nature methods 2017;14(11):1083–1086.

Baron, M., et al. A Single-Cell Transcriptomic Map of the Human and Mouse Pancreas Reveals Inter- and Intra-cell Population Structure. Cell systems 2016;3(4):346–360.e344.

Blondel, V.D., et al. Fast unfolding of communities in large networks. Journal of Statistical Mechanics: Theory and Experiment 2008;2008(10):P10008–10012.

Butler, A., et al. Integrating single-cell transcriptomic data across different conditions, technologies, and species. Nature biotechnology 2018;36(5):411–420.

Cao, Y., Wang, X. and Peng, G. SCSA: A Cell Type Annotation Tool for Single-Cell RNA-seq Data. Frontiers in genetics 2020;11:490–490.

Cortal, A., et al. Gene signature extraction and cell identity recognition at the single-cell level with Cell-ID. Nature Biotechnology 2021.

Cui, Y., et al. Single-Cell Transcriptome Analysis Maps the Developmental Track of the Human Heart. Cell reports (Cambridge) 2019;26(7):1934–1950.e1935.

Ding, J., et al. Systematic comparison of single-cell and single-nucleus RNA-sequencing methods. Nature biotechnology 2020;38(6):737–746.

Efroni, I., et al. Quantification of cell identity from single-cell gene expression profiles. Genome biology 2015;16(1):9–9.

Franzén, O., Gan, L.-M. and Björkegren, J.L.M. PanglaoDB: a web server for exploration of mouse and human single-cell RNA sequencing data. Database : the journal of biological databases and curation 2019;2019.

Grün, D., et al. De Novo Prediction of Stem Cell Identity using Single-Cell Transcriptome Data. Cell stem cell 2016;19(2):266–277.

Kuppe, C., et al. Decoding myofibroblast origins in human kidney fibrosis. Nature 2021;589(7841):281–286.

Lähnemann, D., et al. Eleven grand challenges in single-cell data science. Genome biology 2020;21(1):31–35.

Lawlor, N., et al. Single-cell transcriptomes identify human islet cell signatures and reveal cell-type– specific expression changes in type 2 diabetes. Genome research 2017;27(2):208–222.

Mancarci, B.O., et al. Cross-Laboratory Analysis of Brain Cell Type Transcriptomes with Applications to Interpretation of Bulk Tissue Data. eNeuro 2017;4(6):ENEURO.0212-0217.2017.

Miao, Z., et al. Putative cell type discovery from single-cell gene expression data. Nature methods 2020;17(6):621–628.

Muraro, Mauro J., et al. A Single-Cell Transcriptome Atlas of the Human Pancreas. Cell systems 2016;3(4):385–394.e383.

Pliner, H.A., Shendure, J. and Trapnell, C. Supervised classification enables rapid annotation of cell atlases. Nature methods 2019;16(10):983–986.

Segerstolpe, Å., et al. Single-Cell Transcriptome Profiling of Human Pancreatic Islets in Health and Type 2 Diabetes. Cell metabolism 2016;24(4):593–607.

Shao, X., et al. scCATCH: Automatic Annotation on Cell Types of Clusters from Single-Cell RNA Sequencing Data. iScience 2020;23(3):100882.

Tian, L., et al. Benchmarking single cell RNA-sequencing analysis pipelines using mixture control experiments. Nature methods 2019;16(6):479–487.

Tucker, N.R., et al. Transcriptional and Cellular Diversity of the Human Heart. Circulation (New York, N.Y.) 2020.

Vieira Braga, F.A., et al. A cellular census of human lungs identifies novel cell states in health and in asthma. Nature medicine 2019;25(7):1153–1163.

Wang, L., et al. Single-cell reconstruction of the adult human heart during heart failure and recovery reveals the cellular landscape underlying cardiac function. Nature cell biology 2020;22(1):108–119.

Wolf, F.A., Angerer, P. and Theis, F.J. SCANPY: large-scale single-cell gene expression data analysis. Genome biology 2018;19(1):15–15.

Zhang, A.W., et al. Probabilistic cell-type assignment of single-cell RNA-seq for tumor microenvironment profiling. Nature methods 2019;16(10):1007–1015.

Zhang, X., et al. CellMarker: a manually curated resource of cell markers in human and mouse. Nucleic acids research 2019;47(D1):D721–D728.

Zhang, Z., et al. SCINA: A Semi-Supervised Subtyping Algorithm of Single Cells and Bulk Samples. Genes 2019;10(7):531.

Zheng, G.X.Y., et al. Massively parallel digital transcriptional profiling of single cells. Nature communications 2017;8(1):14049–14049.

