## Supplementary Figures and Tables for "MACA: Marker-based automatic cell-type annotation for single cell expression data"

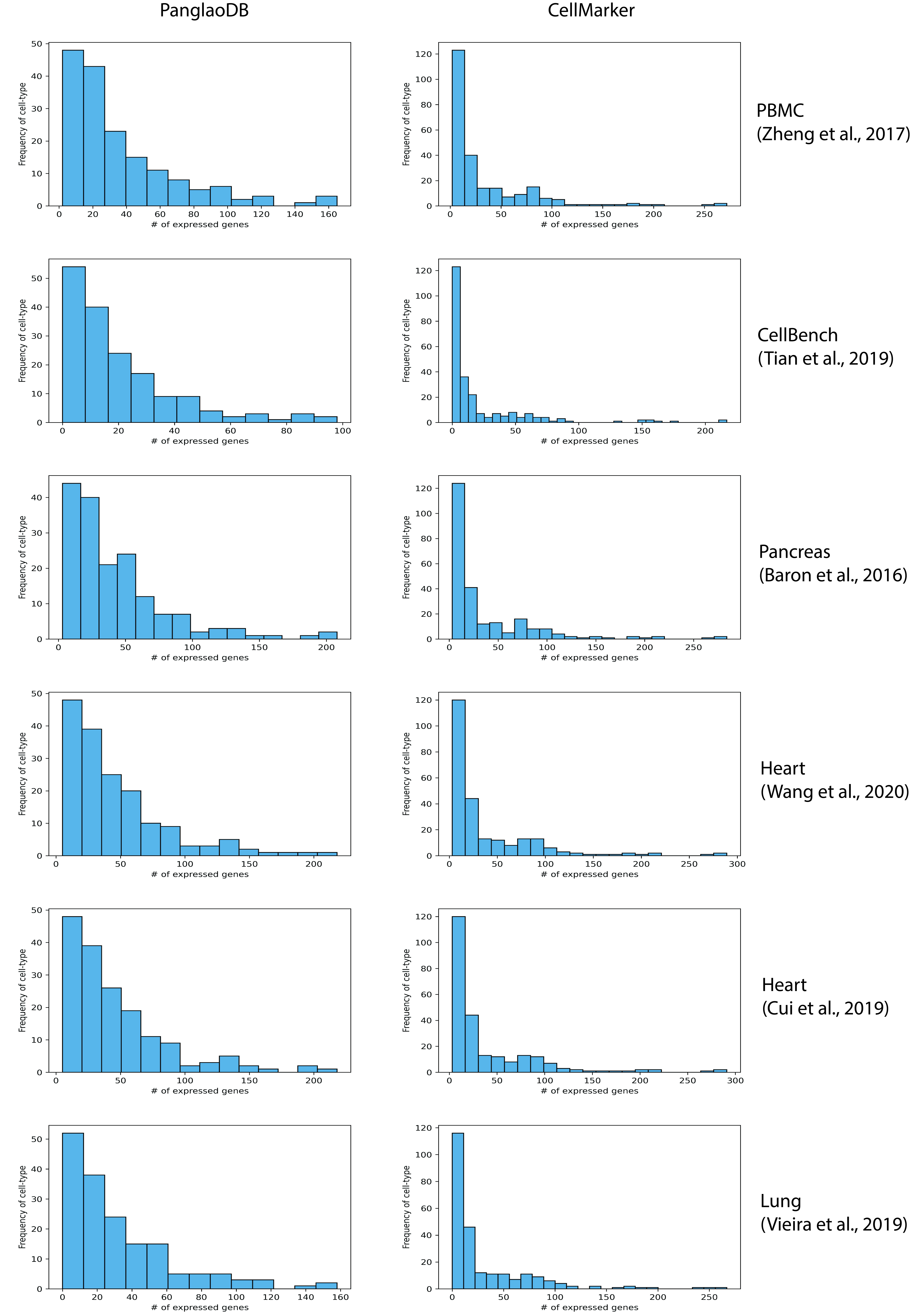
**Supplementary Figure S1**

**Supplementary Figure S1**. Expressed marker genes of cell-types in PanglaoDB and CellMarker across 6 single cell datasets. Any marker genes in either PanglaoDB or CellMarker would be regarded as expressed genes if they have non-zero values in at least one cell. Histograms show the number of expressed marker genes against the number of cell-types.

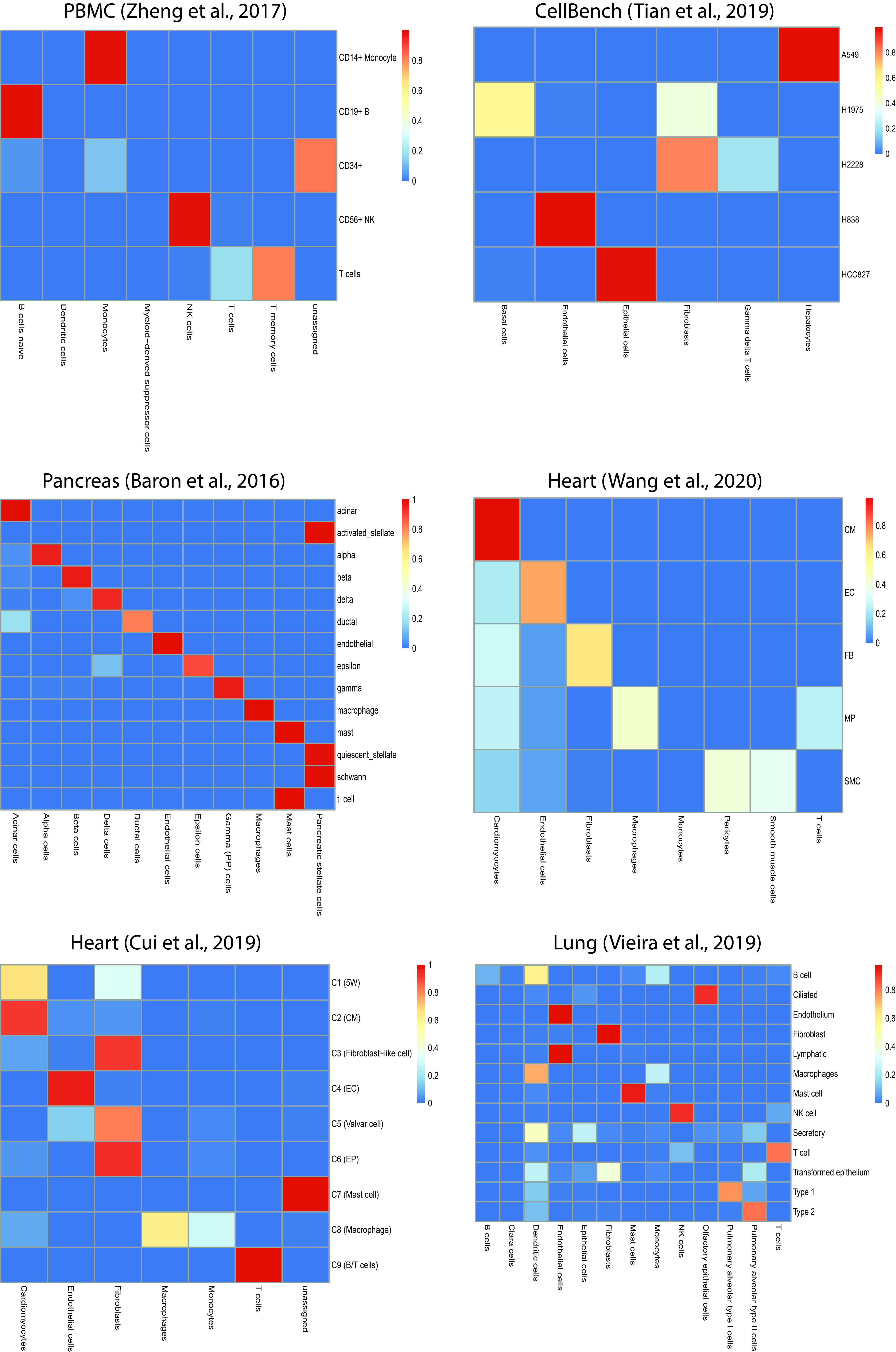
**Supplementary Figure S2**

**Supplementary Figure S2**. Normalized confusion matrices between authors’ annotation on y-axis and annotation by MACA on x-axis, for 6 single cell data used in this study. Each row stands for cell-type label in authors’ annotation, and each column is cell-type label in MACA’s annotation. Confusion matrix was normalized by dividing the sum of each row. A high value means that MACA annotation agrees well with authors’ annotation.

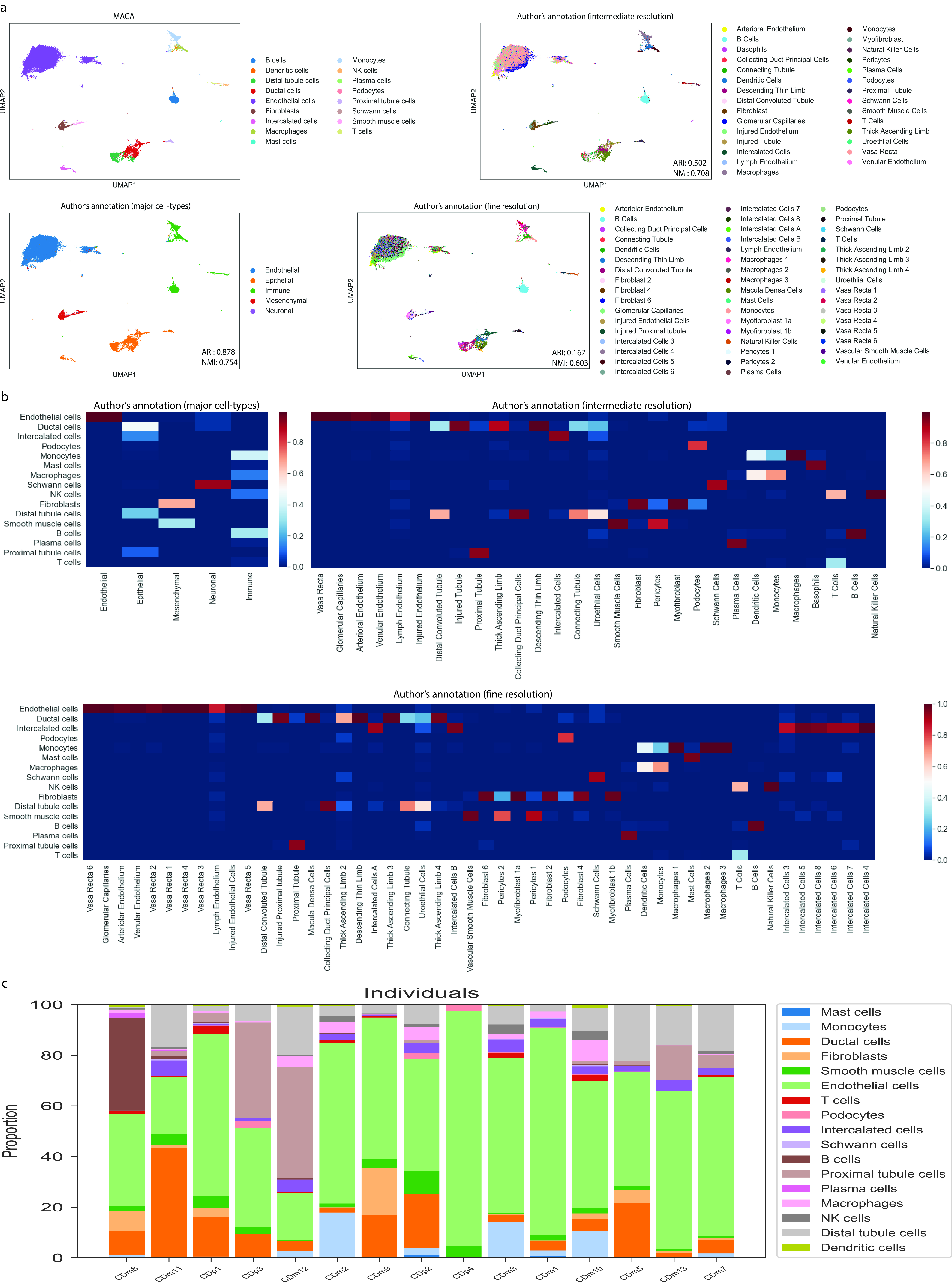
**Supplementary Figure S3**

**Supplementary Figure S3**. Integrated annotation of human kidney (CD10^-^). a, MACA’s annotation is shown in the top. Author reported 3 levels of annotations, with 5 major cell-types, 29 intermediate cell-types, and 50 fine cell-types. MACA’s annotation is compared to all 3 resolutions by ARI and NMI. b, Confusion matrices between MACA’s annotation and 3 author’s annotations. Each column stands for cell-type label in authors’ annotation, and each row is cell-type label in MACA’s annotation. Confusion matrix was normalized by dividing the sum of each column. c, Cellular component analysis across different individuals.

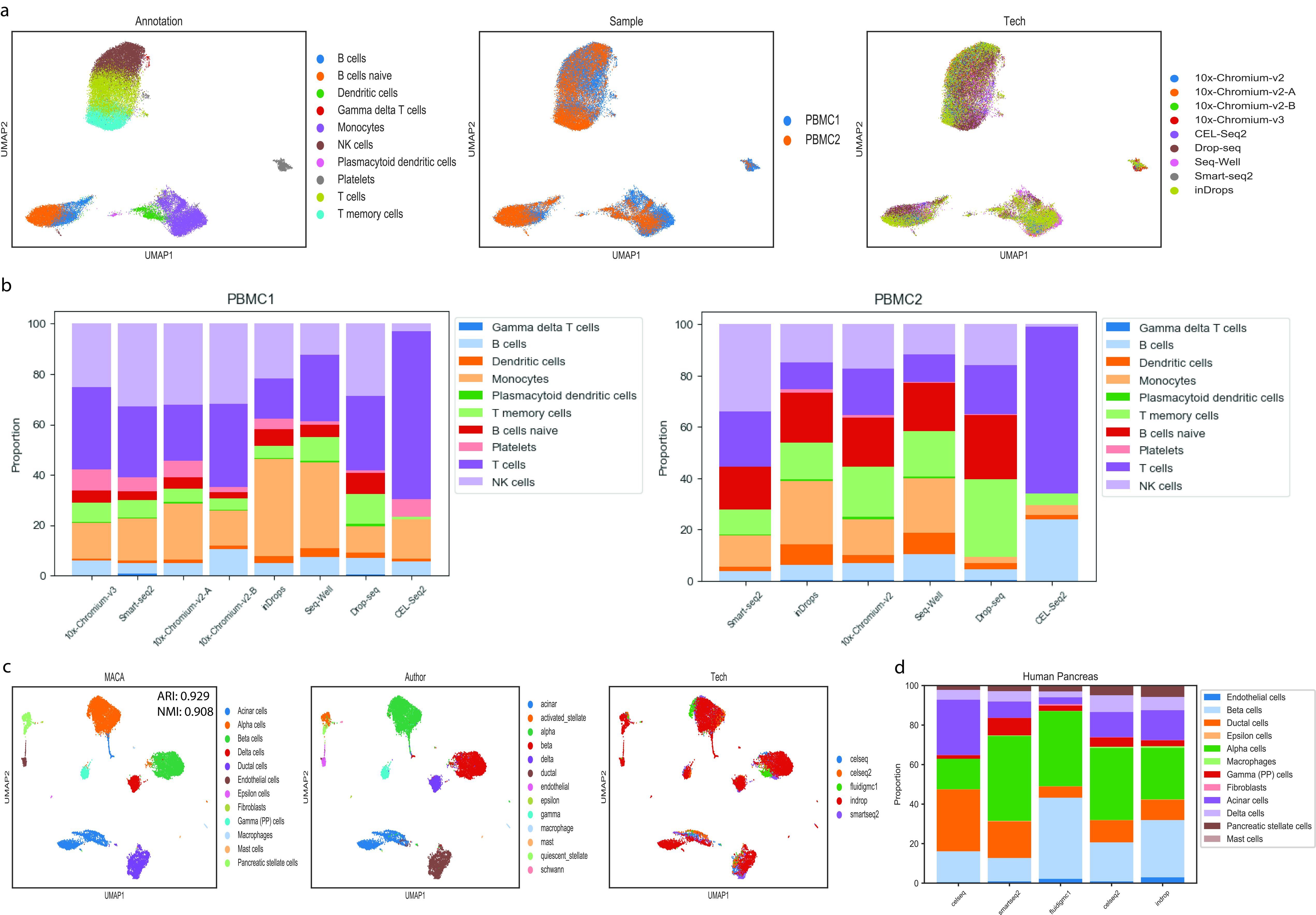
**Supplementary Figure S4**

**Supplementary Figure S4**. Integrated annotation of human PBMC and pancreas data across different single cell platforms. a, UMAP visualization of human PBMC data. Cells are colored according to annotation by MACA (left), source of sample (middle), and platform (right). UMAP dimension reduction is based on PlinerScore with PanglaoDB as marker database. b, cellular component analysis of human PBMC data. Proportion of each cell-type identified by MACA is calculated for each platform for two PBMC samples, separately. c, UMAP visualization of human pancreas data. Cells are colored according to annotation by MACA (left), author-reported cell-types (middle), and platform (right). UMAP dimension reduction is based on PlinerScore with PanglaoDB as marker database. d, cellular component analysis of human pancreas data. Proportion of each cell-type identified by MACA is calculated for each platform.

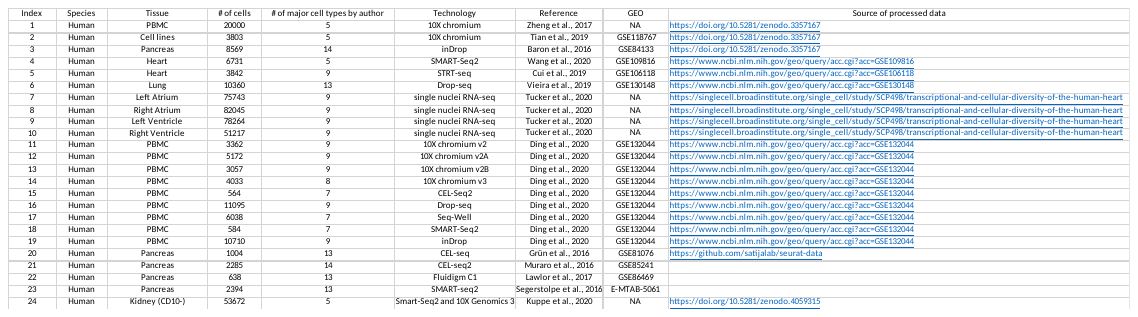
**Table S1**

**Supplementary Table S1.** Summary of datasets used in this study. MACA was developed and its parameters for choosing the optimal scoring and underlying marker database were optimized on 6 datasets (PBMC, CellBench, Pancreas, Heart and Lung). Platform validation was conducted on human PBMC data across 9 single cell RNA-seq platforms and human pancreas data across 5 platforms. Furthermore, MACA’s annotation resolution is compared with 3 different annotation resolutions by author in human kidney (CD10-) data. Fully optimized MACA was then tried on the human 4 chamber heart dataset obtained from Tucker et al., 2020.

**Table S2**

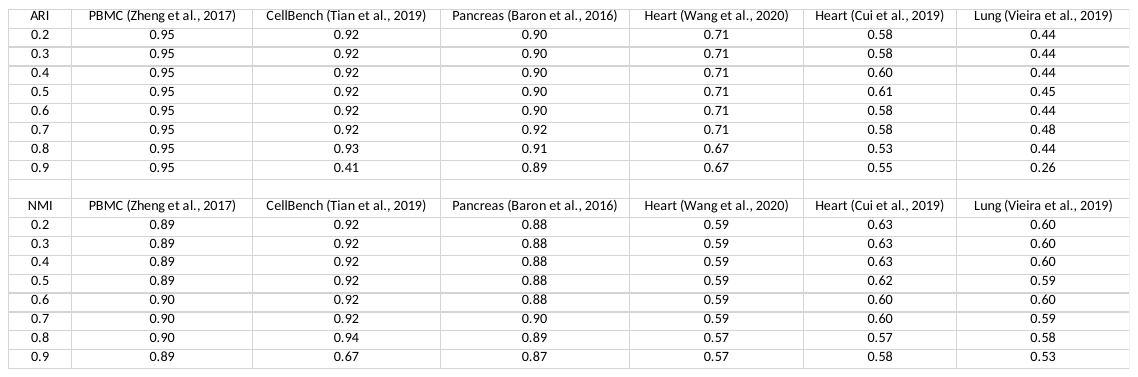

**Supplementary Table S2.** Evaluation of threshold of consensus between Label 1 and Label 2. The first column is the threshold used. Clusters in Label 2 will be assigned a cell-type label in Label 1, only when the proportion of cells in that cluster sharing the same cell-type label is larger than the threshold.

**Table S3**

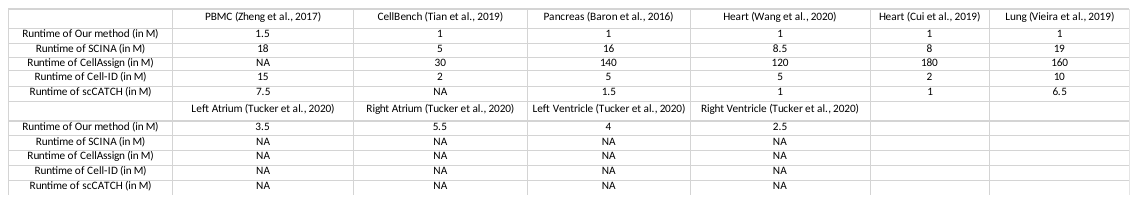

**Supplementary Table S3.** Runtime of 5 annotation tools across 10 datasets. MACA was much shorter than other methods tested in this benchmark, running a workstation with 16-core CPU and 64GB memory.

**Table S4**

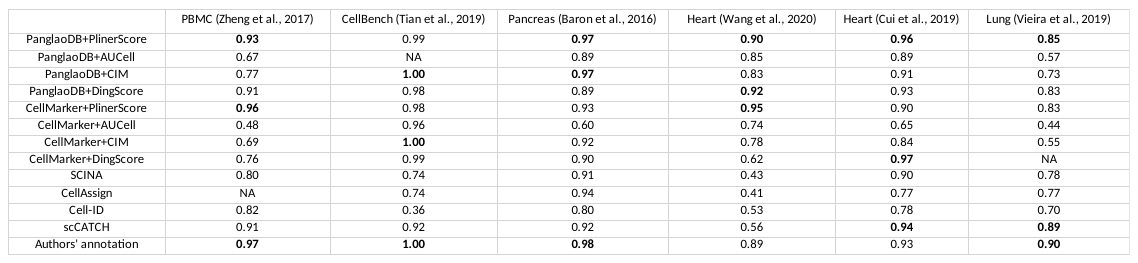

**Supplementary Table S4.** Mean accuracy of 5-fold SVM classifier. SVM classifiers were trained with labels from original reports, or generated by MACA, CellAssign, SCINA, Cell-ID, and scCATCH. Each data was split into 5 folds. Classifiers were trained on 4 folds, and they used the rest 1-fold to report accuracy. Results here came from the mean accuracy of 5-fold training. The highest accuracies obtained for that dataset is shown in bold. For most datasets, PanglaoDB+PlinerScore and authors’ annotations achieve the highest accuracy in SVM classification.

**Table S5**

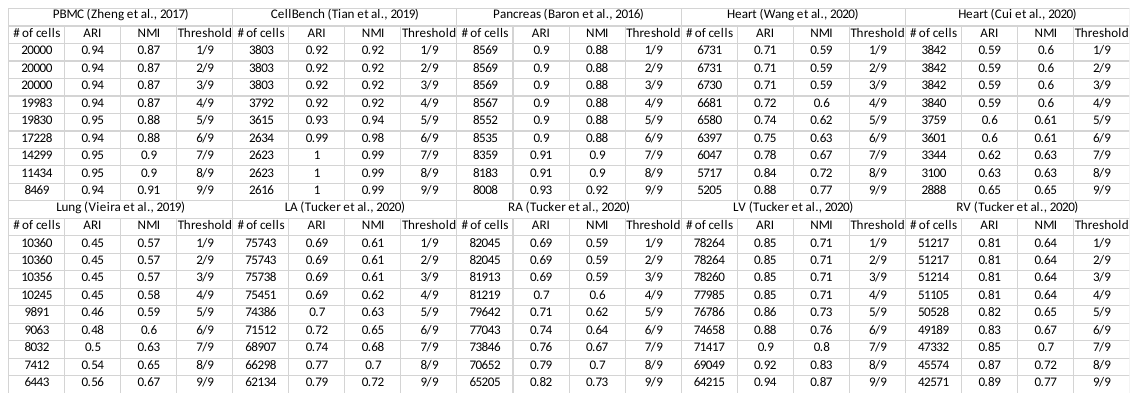

**Supplementary Table S5.** ARI/NMI against the number of cells to be annotated when different thresholds are used. In this study, we performed 9 different settings of clustering to get 9 different cell-type annotations. MACA adopts voting for ensemble annotation. Cells are finally annotated as the cell-type that gets the greatest number of votes. Increasing threshold of voting (the number of votes) to remove cells with less consistent votes inevitably reduces the number of cells to be included in the final annotation. However, comparing annotations of rest of cells by MACA shows increasing agreement with authors’ annotations (increases of ARI and NMI).

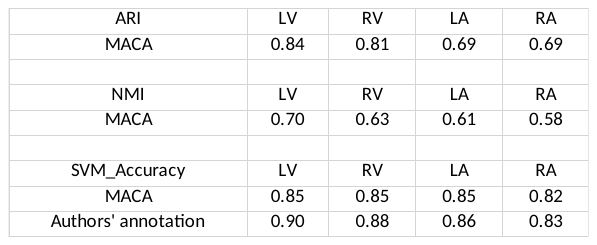
**Table S6**

**Supplementary Table S6.** Performance of MACA in 4 human-chamber single-nuclei RNA-seq datasets, measured by ARI, NMI, and Accuracy of SVM classifier. RA: right atrium; LA: left atrium; RV: right ventricle; LV: left ventricle. The final MACA setting is using PanglaoDB as marker reference and PlinerScore as cell-type scoring method. The performance in human 4 chamber data (Tucker et al., 2020) is quantified by ARI and NMI against author’s annotations (top 4 rows). Both MACA’s and author’s annotation were used to train SVM classifiers. Datasets were split into 5 folds. SVM classifiers were trained on 4-fold data and tested on the rest 1-fold. Means of accuracies were reported to show how reasonable MACA’s and authors’ annotations are (bottom 2 rows).

**Table S7**

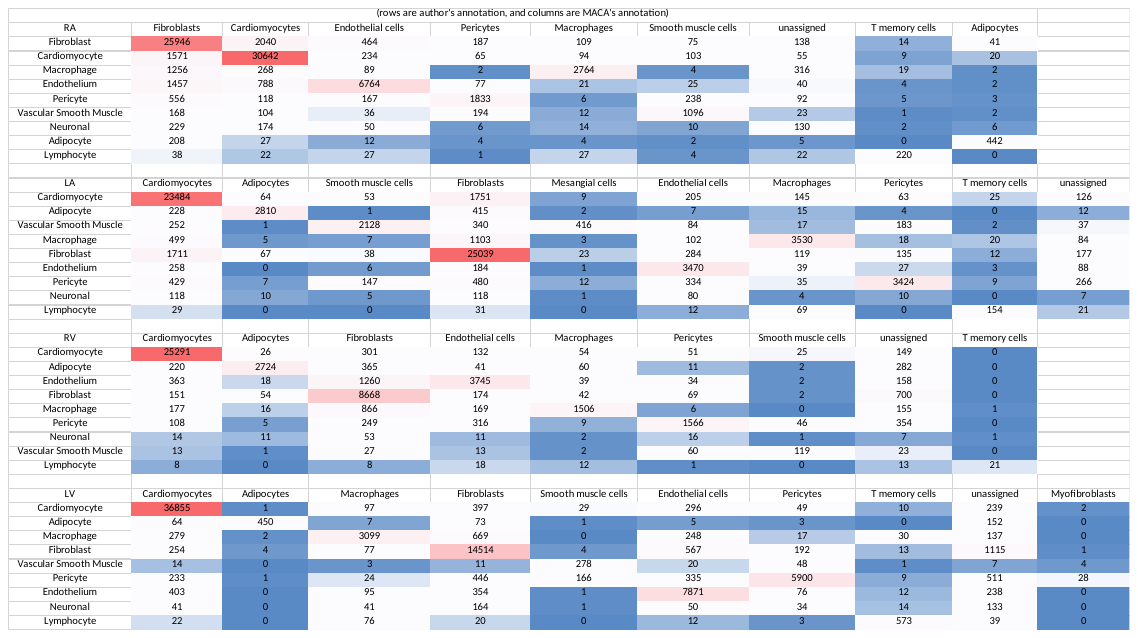

**Supplementary Table S7a.** Confusion matrix between author-reported label and MACA’s annotation in 4 human-chamber sing-nuclei RNA-seq datasets. RA: right atrium; LA: left atrium; RV: right ventricle; LV: left ventricle. Each row represents the number of cells annotated by author, and each column is MACA’s annotation.

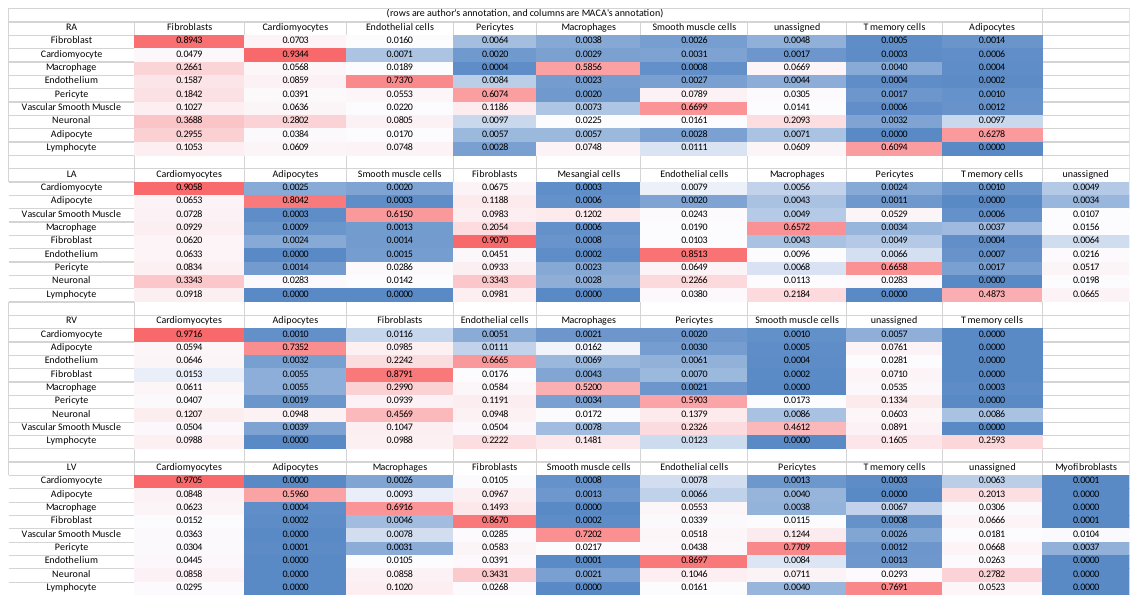

**Supplementary Table S7b.** Confusion matrix between author-reported label and MACA’s annotation in 4 human-chamber sing-nuclei RNA-seq datasets. RA: right atrium; LA: left atrium; RV: right ventricle; LV: left ventricle. From Supplementary Table S7a, each entry is divided by the sum of row, which the total number of cells of each cell-type by author’s annotation. Supplementary Table S7b shows percentages of each cell-type in author’s annotation but in all other cell-types in MACA’s annotation.
